# Radiographic evaluation of subcutaneously injected, water-soluble, iodinated contrast for lymphography

**DOI:** 10.1101/325183

**Authors:** Christopher E. Lee, Brad M. Matz, Robert C. Cole, Harry W. Boothe, D Michael Tillson

## Abstract

Sentinel lymph node (SLN) mapping is common in many types of human cancers, and is gaining utility in veterinary medicine. There are currently many different methods described in veterinary medicine for pre-operative SLN mapping, however, most of these are restricted to referral institutions due to cost and need for specialized equipment. The purpose of this prospective, pilot study was to evaluate the feasibility of radiographic evaluation of water-soluble, iodinated contrast (WIC) injected subcutaneously for lymphography in dogs. Eight dogs were injected with 1-2 milliliters of WIC into the subcutaneous tissues overlying the tarsus in 4 separate locations mimicking a circumferential, peri-tumoral injection. Radiographs were taken at select time points up to 50 minutes. Image sequences were evaluated by a single, board-certified radiologist. All 8 dogs had visible contrast-enhancing lymphatic channels. Median time to lymphatic enhancement was immediately post-injection. Seven dogs (88%) had 8 contrast enhancing lymph nodes (7 popliteal and 1 superficial inguinal). Median time to lymph node enhancement was 20 minutes. In this study, the plantar aspect of the pes drained to the superficial inguinal lymph node, and the dorsal aspect of the pes drained to the popliteal lymph node. Subcutaneously-injected WIC was readily identifiable in the lymphatic channels and draining lymph node(s). Subcutaneously injected WIC may offer a practical alternative to previously described pre-operative methods of SLN mapping. Additionally, one cannot assume that the popliteal lymph node alone, drains the distal pelvic limb.

## Introduction

Sentinel lymph nodes (SLN) are defined as the first lymph node(s) receiving lymphatic drainage from a tumor. Sentinel lymph node mapping and biopsy is performed to stage many types of human cancers[1–4] and is gaining increasing utility in veterinary medicine.[5–13] Sentinel lymph nodes can be presumed based on location of a primary lesion; however, a recent study showed that 8 of 20 dogs (40%) with naturally occurring mast cell tumors demonstrated aberrant lymphatic drainage from that expected based on anatomic location.[14] Additionally, one veterinary study showed that evaluation of only the mandibular lymph nodes could result in up to a 45% under-diagnosis of metastases of oral tumors.[15] Several studies have shown that lymph node metastasis has prognostic implications in different types of cancer, and removal of lymph nodes with confirmed metastasis improves survival in both human [3, 4, 16, 17] and veterinary medicine.[14, 18–25] Based on this information, the importance of identification and sampling of the SLN is clear.

Described methods of pre-operative SLN mapping in veterinary medicine include lymphoscintigraphy,[14] contrast-enhanced ultrasonography,[6, 26] lipid-soluble iodinated contrast (LIC) with radiography^12,14,15^ or computed tomography,[5, 7, 8, 27, 28] and water-soluble iodinated contrast (WIC) with computed tomography.[13, 28–30] Lymphoscintigraphy requires special licensure and specialized equipment that limit the availability of this diagnostic technique. Similarly, the available literature only describes the use of WIC with computed tomography for lymphography. This requires referral to specialty practices or academic institutions in most instances, which may preclude access to this diagnostic technique. While LIC can be used for radiographic evaluation and computed tomographic evaluation, LIC is expensive and can be difficult to obtain. Additionally, optimal imaging evaluation does not occur until 24 to 48 hours after administration[5,7] which adds additional cost and time.

Previous studies have shown that subcutaneously administered WIC is rapidly absorbed into the local lymphatic channels.[8, 9, 11, 12, 28, 30, 31] However, such studies only describe evaluating SLN and lymphatic channels using computed tomography. Development of a more economic and readily available technique for SLN mapping could be beneficial to veterinary oncology patients. Water-soluble contrast is inexpensive and available in most veterinary practices, including primary care veterinary facilities. Radiography is also more readily available in veterinary practices than is computed tomography.

The objective of this study was to identify the feasibility of lymphography in healthy dogs using iopamidol (IS0VUE-370, Bracco, Milan, Italy), a WIC, using digital radiography. The hypotheses of this study were that subcutaneously-injected WIC would be identifiable and traceable in the lymphatics and the draining lymph node. Additionally, it was hypothesized that the popliteal lymph node would be the primary lymphatic drainage of the distal pelvic limb.

## Materials and Methods

A prospective, pilot study was designed. The study protocol was approved by the Institutional Animal Care and Use Committee at Auburn University (Protocol #2017-3199). All dogs underwent a thorough physical examination by one of the investigators (CEL) to rule out any pre-existing conditions. All dogs were re-examined immediately after subcutaneous injection of the WIC, after recovery from sedation, and 24 hours post-injection of WIC to assess adverse events at the injection sites.

All dogs were sedated with dexmedetomidine (Dexdomitor, Zoetis, New Jersey, United States) at 10-15 μg/kg IM and butorphanol (Torbugesic, Zoetis, New Jersey, United States) at 0.3-0.4 mg/kg IM, and an intravenous catheter was placed in a cephalic vein. Dogs were given additional dexmedetomidine intravenously as necessary to obtain appropriate chemical restraint to acquire adequate radiographic images. At the conclusion of the radiographic study, the dexmedetomidine was reversed with an equal volume of atipamezole (Antisedan, Zoetis, New Jersey, United States) administered intramuscularly.

Water-soluble, iodinated contrast (1-2 milliliters) was injected subcutaneously using a 21ga hypodermic needle into 4 separate locations in equal aliquots (0.25-0.5 milliliters/site) around a single point overlying the plantar aspect of the pes at the level of the metatarsals, mimicking the circumferential, peri-tumoral injections used for other types of SLN lymphography[5, 14, 28, 29]. A coin flip was used to determine which pelvic limb, left or right, would be injected and imaged on the first dog. The limb injected and imaged in subsequent dogs was alternated thereafter resulting in 4 left and 4 right limbs being evaluated.

Digital radiographs (Siemens Ysio, Siemens Healthcare, Munich, Germany) were acquired at standardized time intervals including prior to contrast injection, 0, 3, 5, 10, and 20 minutes post-injection to assess the course of the iodinated contrast medium. The kVp varied for each animal based on a radiographic technique chart and the mAs was set at 4.5. The radiographs were mediolateral projections of the entire pelvic limb (left or right) from the pelvis to the digits in all cases. All images were stored in DICOM format on the local PACS. Images were reviewed by one board-certified veterinary radiologist (RCC) on a dedicated DICOM viewer software (eFilm version 3.3, MERGE, WI, USA).

Parameters recorded included initial time to lymphatic channel enhancement, course of the lymphatic channel(s), which lymph node(s) enhanced, if any, and initial time of lymph node enhancement.

## Results

Eight 9-month old, intact male Beagles weighing 13 to 14 kg obtained from a licensed USDA source were included in this study. All animals were deemed healthy on the basis of physical examination. These dogs were obtained for a use unrelated to this study, however, no other interventions were done prior to this study.

Mediolateral radiographic images were obtained of the injected limb pre-injection and at times 0, 3, 5, 10, and 20 minutes post-injection in all dogs. Based on degree of lymph node enhancement in the first three dogs the study protocol was modified to include radiographs up to 60 minutes if necessary to achieve lymph node enhancement. Thus, mediolateral images were taken at 30 and 40 minutes in 5 of the 8 dogs and at 50 minutes in 1 dog.

One milliliter of WIC was injected into the subcutaneous tissue on the plantar aspect of the pes in 1 dog, 2 milliliters of contrast was injected on the plantar aspect in 6 dogs, and 1 milliliter was injected into the subcutaneous tissues of both the plantar and dorsal aspects, for a total of 2 milliliters, of the pes in 1 dog. The volume of WIC was increased after the first dog, as the contrast material was visible within the lymphatics and approaching the popliteal lymph node within the study period, however, nodal enhancement did not occur. The final dog in this study was injected on both the plantar and dorsal aspects of the pes in response to identification of a lymphatic pathway from the dorsal aspect of the pes to the popliteal lymph node.

All dogs had visible enhancement of lymphatic channels. The median time to initial enhancement of lymphatic channels was immediately post injection (range 0-5 minutes) and depletion of lymphatic channel enhancement was not identified during the study period.

Seven of 8 dogs had enhancement of multiple lymphatic channels. Three patterns of lymphatic drainage were observed: lymphatic drainage from the plantar aspect of the pes coursed caudally over the tarsus, caudal to the tibia and the stifle, and consistently drained to the region of the superficial inguinal lymph node (n=7); lymphatic drainage from the dorsal aspect of the pes continued cranially over the tarsus and, at the level of the distal third of the tibia, transitioned to a more caudal position in its course to the popliteal lymph node (n=7); and lymphatic drainage via a third lymphatic channel which was neither dorsal nor plantar drained to the popliteal lymph node (n=1) (Fig 1). Six of 8 dogs had both the dorsal and plantar pathways enhance, 1 dog had both the third channel and plantar pathways enhance, and 1 dog had only the dorsal pathway enhance. Lymphatic communication between the dorsal and plantar aspect of the pes at the level of the metatarsals was identified in 3 of 6 dogs with both the dorsal and plantar drainage pathways. The dog with injections of both the dorsal and plantar aspects of the pes occurred was the only dog not to have multiple lymphatic pathways enhance. Only the dorsal pathway enhanced in this dog. Difficulty injecting the contrast into the subcutaneous tissue on the plantar side and leakage from the previous injection sites in this area led to a smaller volume of contrast injected at this location than intended.

**Fig 1.**
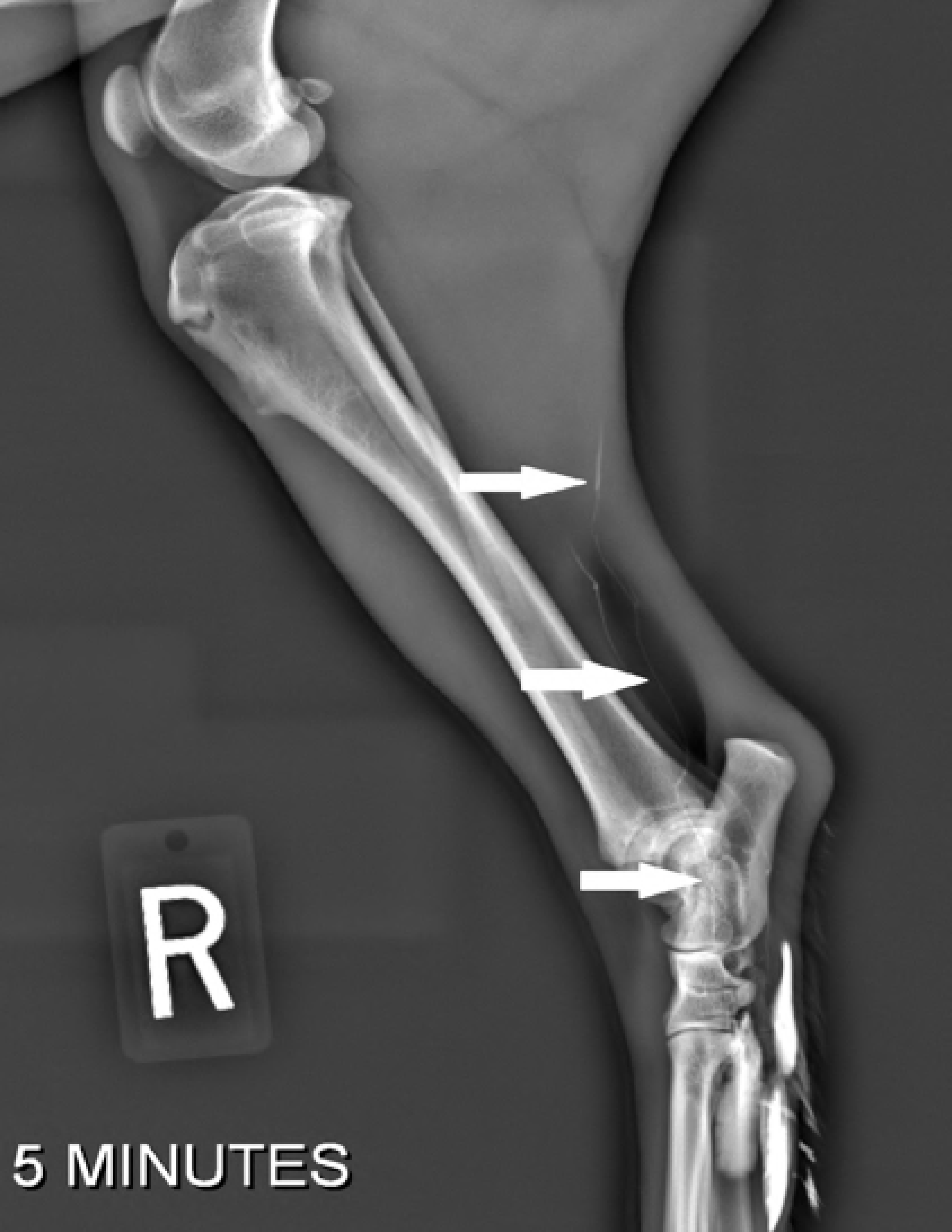
Third lymphatic channel. This mediolateral radiograph of the right pelvic limb depicts the third lymphatic channel, labelled with the long, solid white arrows, identified in Dog 1. Like the dorsal lymphatic channel identified in the other dogs, this channel appeared to terminate in the popliteal lymph node.

Seven of eight dogs (88%) had enhancement of 8 lymph nodes (Fig 2) including 7 popliteal lymph nodes and 1 superficial inguinal lymph node. Median time for initial enhancement of the lymph nodes was 20 minutes (range 5-50 minutes). Median time for maximal enhancement of the lymph nodes was 30 minutes (range 10-50 minutes). In 3 of 7 dogs and 4/8 lymph nodes that enhanced, maximum contrast enhancement occurred at the last radiograph taken. All but 1 dog exhibited enhancement of the draining lymph node(s) that continued to increase or remained static once maximal enhancement was reached throughout the duration of the study period. One lymph node, which had initial enhancement at 5 minutes, had mild decrease of contrast enhancement at 40 minutes, but the lymph node remained enhanced in comparison to precontrast images. One dog had enhancement of the lymphatic vasculature, but not the draining lymph node. This was the initial dog studied injected with only 1 ml of contrast, prompting the increase to 2 ml for subsequent evaluations. Enhancement of both the superficial inguinal and popliteal lymph node occurred in one dog at 20 and 50 minutes, respectively (Fig 3). Enhancement of separate afferent lymphatics leading to each lymph node from the injection site were identifiable, and there were no visible efferent lymphatics from either lymph node.

**Fig 2.**
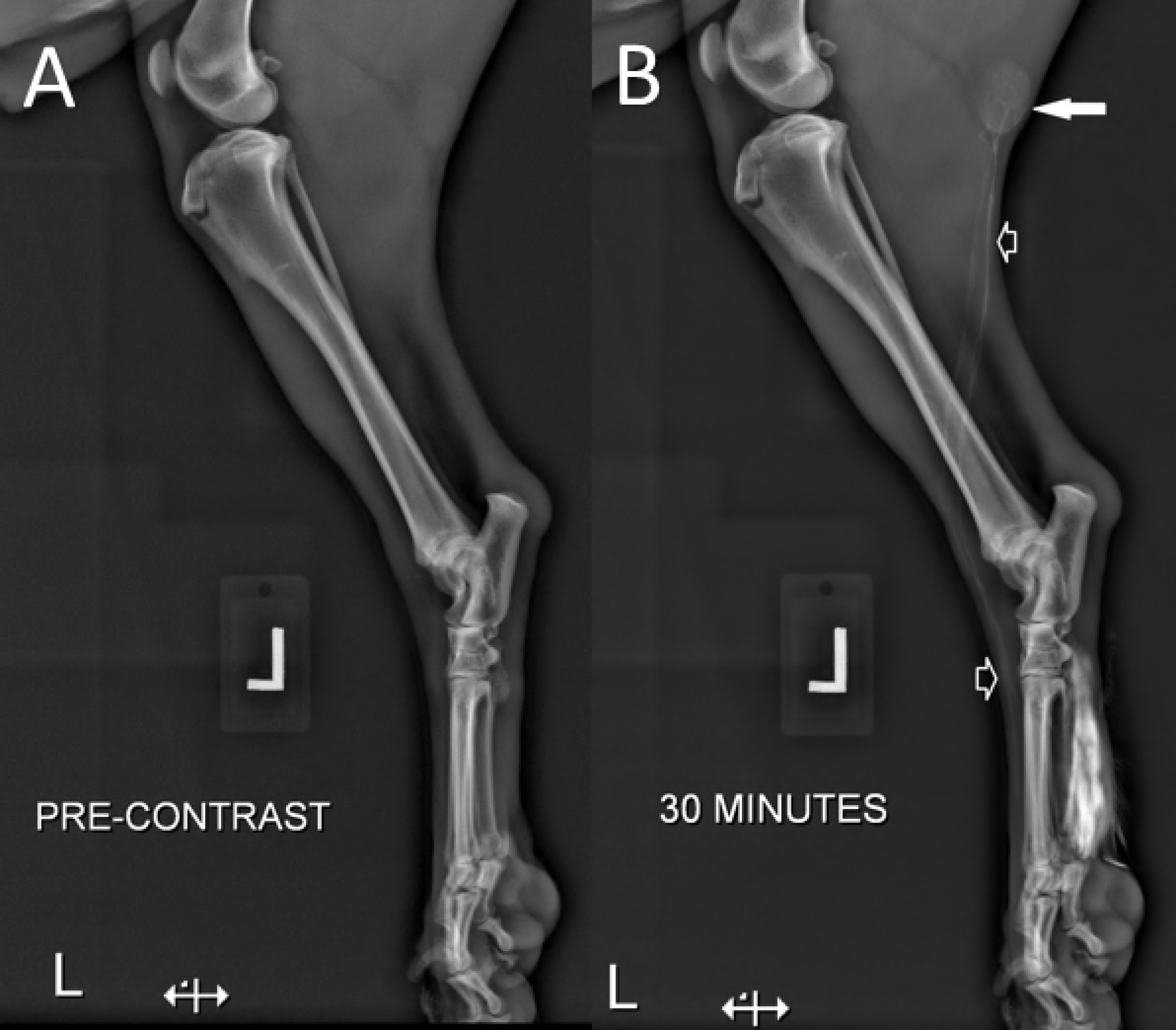
Popliteal lymph node enhancement and dorsal and planter lymphatic channels. Mediolateral radiographs of the left pelvic limb showing the pre-contrast (A) and 30 minutes post-contrast (B) images. In image B, the dorsal lymphatic channel, identified with the short, white outlined arrows, is contrast enhanced and leads to the contrast enhanced popliteal lymph node identified with the long, solid white arrow.

**Fig 3:**
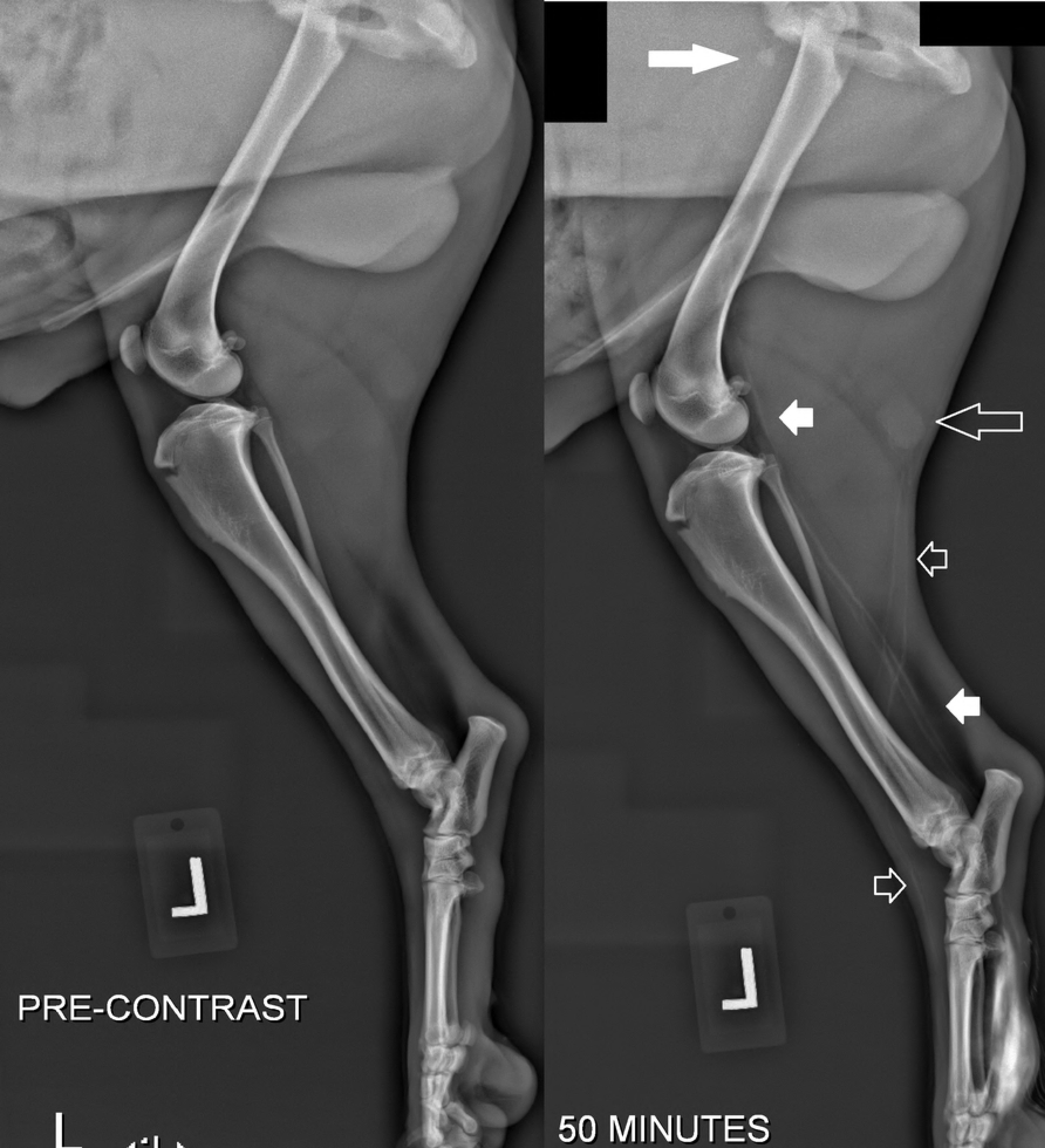
Popliteal and inguinal lymph node enhancement. Mediolateral radiographs of the left pelvic limb showing the pre-contrast (A) and 50 minutes post-contrast (B) images. The dorsal lymphatic channel, depicted by the short, outlined white arrows, is contrast enhanced and leads to the contrast enhanced popliteal lymph node, identified by the long, outlined white arrow. The caudal lymphatic channel, depicted by the short, solid white arrows, is contrast enhanced and identified leading towards the contrast enhanced inguinal lymph node, depicted by the long, solid white arrow.

Adverse events associated with the contrast injection site occurred in all dogs but were deemed minor. No complications requiring additional monitoring or therapy were observed in any of the dogs. There was minimal local hemorrhage immediately after injection. This resolved with application of digital pressure. Leakage of contrast material occurred at previous injection sites as additional sites were injected. One dog required additional dexmedetomidine during injection of the contrast due to stimulation immediately after insertion of the needle in the first injection location. Palpable swelling occurred immediately post injection in all dogs. The swelling dissipated in all dogs by the following day.

## Discussion

To the authors’ knowledge, this is the first study evaluating the effectiveness of radiography for lymphography using subcutaneously-injected WIC. Findings of this study show indirect lymphography with WIC to be feasible for evaluating the lymphatic drainage of the pes.

Three patterns of lymphatic drainage from the pes were identified. Although lymphatic communication between the plantar and dorsal aspects of the pes was only identifiable in 3 of the 6 dogs in which both the dorsal and plantar drainage pathways were evident, it is likely that these communications still exist in the other dogs as the dorsal pathway begins at the level of the metatarsals in all 6 dogs. A third lymphatic channel was identified in 1 dog. It was undetermined whether this channel was medial or lateral because orthogonal images of the limb were not obtained. However, the ultimate course of this pathway was similar to the dorsal lymphatic drainage and terminated in the area of the popliteal lymph node. Obtaining the orthogonal radiographs would not have added particular value to this study, however, if this technique of indirect lymphography is to be used for sentinel lymph node mapping in clinical patients, it is important to get all radiographic images necessary to accurately identify the sentinel lymph node.

All but 1 dog had enhancement of the primary draining lymph node. Although lymph node enhancement was evident within 20 minutes in 6 of the 8 dogs, the authors presume that giving a smaller volume of contrast, may extend the time necessary for contrast enhancement of the draining lymph node. As this was the initial dog of the study receiving 1 milliliter of WIC on the plantar aspect of the pes and having radiographs taken up to 20 minutes post-injection, the role of experimental design on this finding is unclear. In this dog, contrast material was present subjacent to and tracing toward the popliteal lymph node at the 20 minute radiograph, however, it was not identifiable within the lymph node. Including later radiographs in the first study might have improved detection of lymph node enhancement.

Alternatively, 1 milliliter of contrast material may not be sufficient to enhance the draining lymph node in all cases, irrespective of time after injection. In a study by Grimes et al[32], 78% (7/9) of dogs injected with 1 ml of contrast had SLNs identified by computed tomographic evaluation in comparison to 100% (9/9) of dogs injected with 2 milliliters of contrast. Confounding factors theorized to be impacting detection of drainage to the SLN in that study included tumor compression of lymphatics, patient positioning, and/or endotracheal tube ties causing collapse or restriction of lymphatic flow. These factors were not a part of this study suggesting that an additional confounder, such as time or volume of contrast material, was the main factor for the negative result here. Considering these findings, the authors recommend using 2 mls of WIC for lymphographic evaluation with radiographs.

In this study, the achievement of maximal enhancement of the lymph nodes occurred at a median of 30 minutes. However, with the exception of one dog in which lymph node enhancement decreased, maximal lymph node enhancement occurred at the last taken radiograph in 50% of dogs, with the remaining dogs having reached maximal enhancement at the next to last radiograph. It is possible that if the radiographic studies were extended, further enhancement might have occurred. Contrarily, it is also possible that a decrease of contrast enhancement might have occurred in more of these cases if images were acquired at later times due to spread beyond the primary lymph node. There was no observed decrease in lymphatic vasculature enhancement in this study. Therefore, if using this technique for SLN mapping, the authors would recommend obtaining radiographs within 30 minutes post injection to follow the lymphatic channels to the sentinel lymph node and, if necessary, intermittently thereafter until a distinct sentinel lymph node is identifiable.

Limitations of this study included use of young, healthy dogs without masses or lesions whose lymphatic drainage could be mapped and the lack of orthogonal radiographs. However, as the goal of this study was to determine the feasibility of indirect lymphography using WIC and radiographic interpretation. Additionally, the slight variations in WIC injection techniques did not detract from the aforementioned goal. Further studies to assess the utility of this technique for sentinel lymph node mapping in patients with a variety of cutaneous and subcutaneous neoplasms are warranted.

Based on the results of this study, subcutaneously-injected, water-soluble, iodinated contrast material provides a relatively quick and effective means of tracing the lymphatic channels from the pes to the draining lymph node(s). The results of this study also show that one cannot assume that the distal pelvic limb will have primary lymphatic drainage only to the popliteal lymph node.

## Acknowledgements

The authors would like to thank the radiology technicians at Auburn University for their assistance with image acquisition for this project.

